# Programmable Nucleic Acid Sensing in Human Cells Using Circularizable ssDNA

**DOI:** 10.1101/2025.03.12.642784

**Authors:** Ahmed Mahas, Raphael Ferreira, Lisa M. Riedmayr, George M. Church

## Abstract

Programmable technologies that sense specific nucleic acid signatures in living cells and trigger cellular functions hold significant potential for biotechnology and medicine. Here, we developed SONAR (**S**ensing **O**f **N**ucleic acids using **A**SOs and **R**everse-transcriptases), a platform that enables the detection of target DNA and RNA sequences and triggers controlled gene expression in human cells. SONAR operates through circularizable single-stranded DNA (ssDNA) sensors that, upon hybridization with complementary DNA or reverse-transcribed RNA, undergoes target-dependent ligation via endogenous ligases, subsequently driving expression of genetic payloads. For RNA sensing, we employed chemically modified antisense oligonucleotides (ASOs) to prime targeted reverse transcription, generating complementary DNA that promotes ssDNA circularization. We demonstrate SONAR’s ability to detect DNA, exogenous and endogenous RNA transcripts, coupled with a programmable expression of diverse protein payloads, including reporters, recombinases, and genome editors. This platform establishes a versatile framework for targeted nucleic acid detection and inducible gene expression, with broad applications in diagnostics, therapeutics, and synthetic biology.

## Main

Distinct RNA expression profiles and specific DNA mutations are hallmarks of diseases such as cancer, viral infections, and genetic disorders, influencing disease onset, progression, and treatment resistance^1,2^. Detecting these molecular signatures within living cells could greatly enhance precision diagnostics and targeted therapeutics. Biosensors capable of real-time detection would enable earlier disease identification, precise monitoring of treatment responses, and selective therapeutic activation, ultimately improving clinical outcomes. For instance, the detection of specific nucleic acid sequences could trigger programmable outputs such as amplified reporter signals for pathogen detection, the generation of synthetic biomarkers for non-invasive diagnostics, or precise genome editing^3,4^. More advanced approaches could further activate immune responses at tumor or infection sites or initiate apoptotic pathways as kill-switch mechanisms, thereby limiting pathogenic cell proliferation. Importantly, by preferentially activating therapeutic payloads in target cells, these engineered biosensors significantly enhance treatment specificity, thereby reducing off-target effects and associated toxicities.

Recent advancements in nucleic acid-responsive technologies, including eukaryotic toehold switches^5^, circular RNAs^6^, and ADAR-based biosensors^7–9^, represent significant milestones in the field, expanding the capabilities for nucleic acid sensing and controlled cellular responses. However, these approaches rely on intricate setups, complicated designs, are strictly confined to RNA detection, and/or are unable to detect potentially pathogenic single nucleotide polymorphisms (SNPs), ultimately narrowing their potential therapeutic and diagnostic applications. Consequently, there is an unmet need for innovative platforms that overcome these limitations, enabling the detection and integration of diverse nucleic acid signals with high flexibility, sensitivity and specificity, and thus enabling next-generation therapeutic and diagnostic interventions.

While plasmids have traditionally been the dominant platform for nucleic acid-based expression and sensing applications, alternative approaches are expanding the molecular toolkit. Notably, breakthroughs in synthesizing long, high-fidelity single-stranded DNA (ssDNA) molecules have unlocked new opportunities for gene expression^10^, genome editing^11,12^, and advanced synthetic biology applications^13^. Additionally, ssDNA exhibits lower immunogenicity than dsDNA, supporting improved cellular viability and making it more suitable for potential therapeutic applications^11^. Long ssDNA molecules also provide unique structural properties for innovative biosensor designs. Their ability to undergo sequence-specific circularization and ligation enables highly specific nucleic acid sensing within cells.

Building on this concept, we introduce a novel programmable platform, termed SONAR (Sensing Of Nucleic acids using ASOs and Reverse-transcriptases), designed to detect specific DNA and RNA sequences directly within living human cells. SONAR functions by recognizing a targeted complementary nucleic acid sequence, which triggers the circularization of a ssDNA sensor followed by a unique target-dependent ligation mechanism mediated by endogenous cellular ligases. To facilitate RNA detection, antisense oligonucleotides (ASOs) are employed to act as primers within the cells, initiating a targeted reverse transcription using a Moloney murine leukemia virus (MMLV) reverse transcriptase (MMLV-RT). This process produces a complementary reverse-transcribed DNA (RT-DNA), which serves as a “splint” to enable the ligase-mediated circularization of the ssDNA sensor. Once circularized, the ssDNA sensor acts as an efficient, programmable vector, driving the precise expression of genetic payloads within the cell (Fig. 1a).

**Figure 1.**
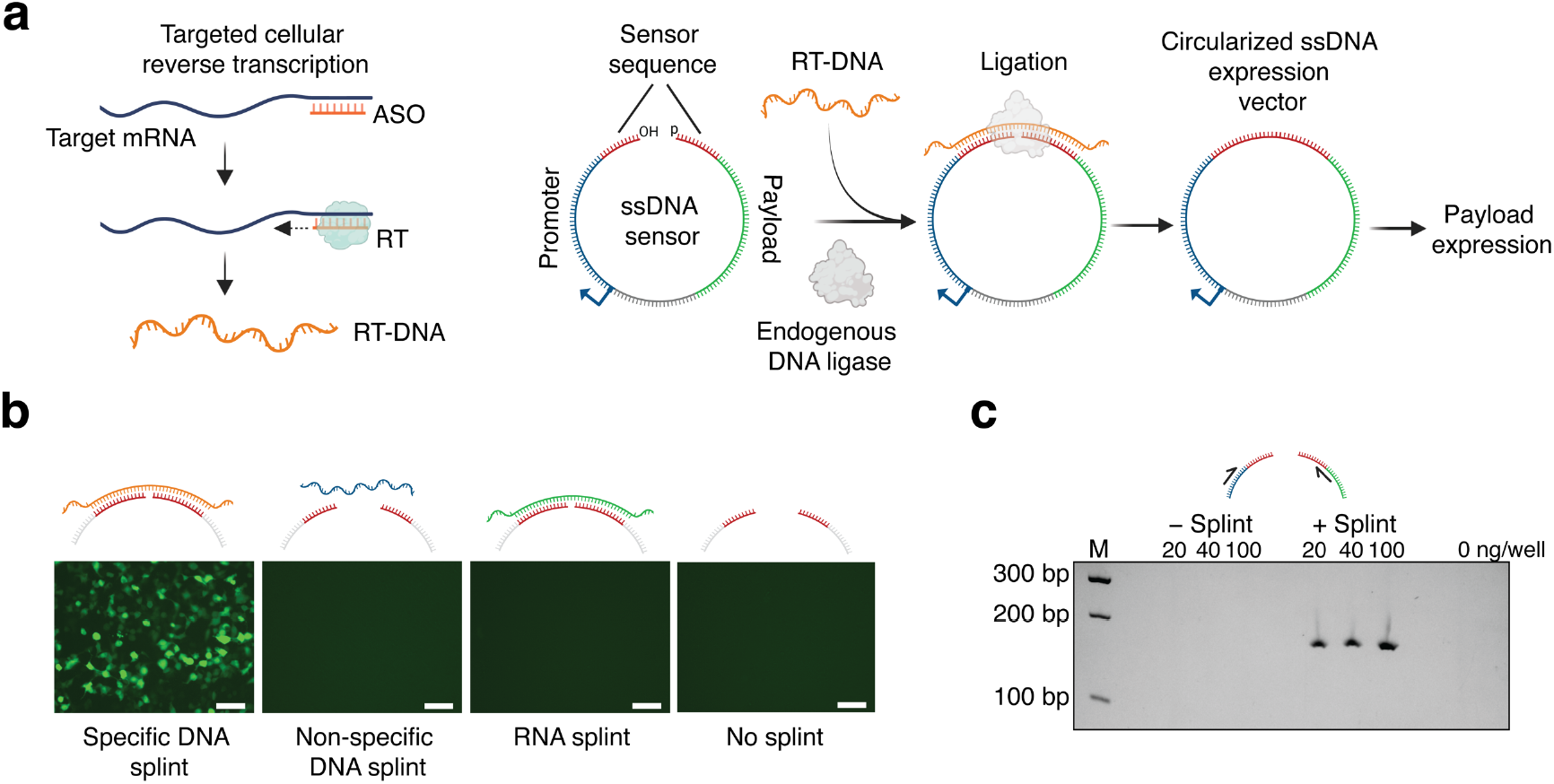
Sensor mechanism and validation. **a**. Schematic illustrating the SONAR design. Antisense oligonucleotides (ASOs) act as primers to mediate targeted reverse transcription of RNA sequences utilizing a reverse transcriptase (RT), producing reverse-transcribed DNA (RT-DNA). The ssDNA sensor contains a sensing sequence complementary to the RT-DNA. Upon hybridizing to the target ssDNA or RT-DNA, ligation-compatible ends form a nicked substrate. Endogenous ligases then circularize the ssDNA sensor, creating a functional expression vector for the payload. **b**. Fluorescence images of HEK293T cells transfected with the ssDNA sensor and either a complementary ssDNA splint or control conditions. Scale bar, 125 µm. **c**. Gel electrophoresis of PCR-amplified sensor from cells transfected with varying sensor concentrations in the presence or absence of a complementary ssDNA splint.

First, we tested whether engineered ssDNA sequences can sense and specifically respond to complementary DNA sequences in human cells. To achieve this, we designed a linear ssDNA sensor consisting of a Green Fluorescent Protein (GFP) reporter, a transcription termination sequence, and a cytomegalovirus (CMV) promoter, arranged in the specified order. In its linear form, the CMV promoter and GFP coding sequence remain physically separated, preventing GFP expression. However, the sensor is phosphorylated at its 5′ end, enabling ligation and circularization when hybridized with a complementary DNA splint. Upon recognition of such a splint, the ligation restores the correct arrangement of the expression cassette, allowing GFP transcription to be initiated. We observed GFP expression 24 hours after co-transfection of the ssDNA sensor with its complementary DNA splint, demonstrating efficient target-dependent activation (Fig. 1b and Supplementary Fig. 1). In contrast, co-transfection with a complementary RNA splint failed to induce GFP expression, confirming the DNA-specific nature of the ligation mechanism. To confirm successful sensor circularization in cells, we amplified the ligation site via PCR using junction-spanning primers. Specific amplification occurred exclusively in cells transfected with both the ssDNA sensor and a complementary DNA splint (Fig. 1c and Supplementary Fig. 2a). Additionally, treatment of the ssDNA sensor with a phosphatase prior to transfection reduced GFP expression, supporting an endogenous ligase-mediated mechanism and highlighting the requirement for a 5′ phosphate for successful circularization (Supplementary Fig. 2b).

Next, we examined the sensor’s capability to detect specific RNA sequences within living human cells. We hypothesized that targeted intracellular reverse transcription of RNA sequences into complementary DNA (RT-DNA) would enable interaction with the sensor, leading to its circularization and payload expression. Previous findings from prime editing studies have demonstrated that targeted reverse transcription in live cells is feasible^14^. To be able to initiate targeted reverse transcription, we designed chemically stabilized ASOs that hybridize to their RNA targets, creating DNA:RNA duplexes. These duplexes then recruit an exogenously supplied reverse transcriptase to mediate a targeted synthesis of complementary DNA (Fig. 1a). For our study, the ASOs were modified with locked nucleic acids (LNAs) throughout their backbone, leaving the terminal four nucleotides unmodified to function effectively as DNA primers for reverse transcription (Supplementary Fig. 2c).

To validate this approach, we designed ASOs targeting exons of two endogenous transcripts, *GAPDH* (glyceraldehyde-3-phosphate dehydrogenase) and *ACTB* (β-actin). HEK293T cells were transfected with reverse transcriptase alone, ASOs alone, or a combination of both. To validate a successful ASO-guided reverse transcription and complementary RT-DNA synthesis, we designed quantitative PCR (qPCR) primers spanning exon-exon junctions. Notably, qPCR as well as gel electrophoresis confirmed precise RT-DNA synthesis only in cells co-transfected with both ASOs and a reverse transcriptase, verifying the specificity and effectiveness of the ASO-guided reverse transcription process (Fig. 2a and Supplementary Fig. 2d).

**Figure 2.**
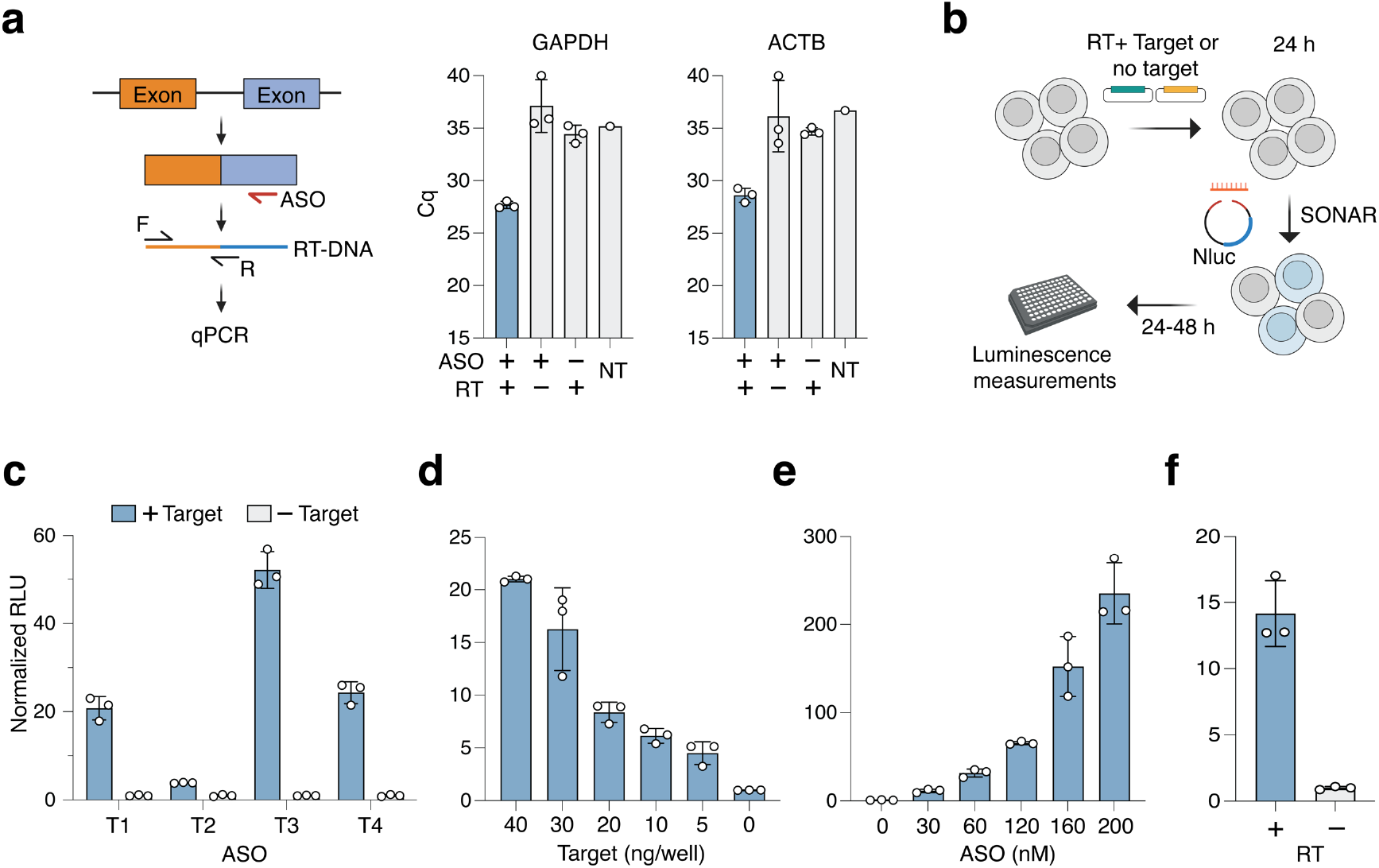
Characterization of SONAR for RNA detection in human cells. **a**. (Left) Schematic of the qPCR assay to analyze the targeted in-cell RT-DNA synthesis. (Right) qPCR analysis of RT-DNA synthesis from two endogenous housekeeping genes. NT: No-template control. **b**. Schematic of sensor detection of target RNA in cells. **c**. Quantification of sensor efficiency in detecting target RNA in human cells. Different ASOs (T1–T4) were tested using plasmids expressing the target sequence (+ Target) or the control plasmid (− Target). For each ASO, the data are presented as fold change relative to the corresponding − target control. **d**. Quantification of sensor efficiency in detecting different target RNA concentrations in human cells with ASO T3. Data are presented as fold change relative to the 0 ng/well target control. **e**. Impact of ASO concentration on sensor efficiency in detecting the target RNA. Data are presented as fold change relative to the 0 nM ASO control. **f**. Effect of reverse transcriptase presence on sensor performance. Data are presented as fold change relative to the no RT control. In **a, c, d, e** and **f**, data represent the mean of *n*=3 biological replicates.

After demonstrating that DNA splints activate the ssDNA sensor and that targeted intracellular reverse transcription can generate complementary RT-DNA from RNA templates, we tested whether these two capabilities can be combined to effectively detect target RNA sequences within living cells. First, we tested whether the sensor could detect exogenous RNA expressed from a plasmid. To achieve this, we designed four distinct ASOs targeting different regions of the target RNA, alongside a control plasmid lacking the target RNA sequence. Cells were co-transfected with a plasmid expressing the MMLV-RT and either the plasmid encoding the RNA target or the control plasmid. After 24 hours, an ASO was co-delivered with the ssDNA sensor encoding a NanoLuc luciferase (Nluc^15^) as a reporter payload (Fig. 2b). All tested ASOs successfully activated the sensor, with the most effective ASO resulting in an approximately 52-fold higher signal than the no-target control (Fig. 2c and Supplementary Fig. 3a). The sensor’s activity increased proportionally with the concentration of the target RNA, demonstrating its ability to quantitatively sense RNA levels in cells (Fig. 2d and Supplementary Fig. 3b). Additionally, increasing ASO concentrations led to enhanced sensor activation, confirming that ASO abundance directly influences detection efficiency (Fig. 2e and Supplementary Fig. 3c).

We also evaluated the influence of nuclear localization signals (NLS) on reverse transcriptase activity, comparing RT variants with and without NLS. RTs without NLS showed a significantly higher sensor activation compared to RTs with NLS, indicating that cytoplasmic reverse transcription is more efficient for this application (Supplementary Fig. 4a). Notably, the absence of an RT completely abolished sensor activation, confirming that reverse transcription is essential for RNA sensing (Fig. 2f and Supplementary Fig. 4a). In addition, we evaluated whether a truncated RT variant lacking the RNase H domain, previously shown to support efficient prime editing in cells^16^, could similarly activate the ssDNA sensor. Both full-length and truncated RT exhibited comparable performance (Supplementary Fig. 4b); thus, the full-length RT without NLS was used in subsequent experiments.

We also explored additional design parameters to optimize our ssDNA sensor system. In our initial designs, we selected 60-nucleotide sensing sequences based on the hypothesis that DNA:DNA hybridization at this length would be inherently stable and efficient. To evaluate the impact of sensing region length, we next compared sequences ranging from 45 to 75 nucleotides. All variants demonstrated comparable levels of activation, indicating that sensing region length has a minimal impact on overall performance within this range (Supplementary Fig. 4c). We also compared alternative sensor configurations by placing the sensing sequence between the stop codon and polyA terminator instead of between the promoter and start codon. While this alternative design was effectively activated upon splint-mediated circularization, our original design demonstrated greater specificity and more robust activation (Supplementary Fig. 4d). Therefore, we chose the original sensor configuration and the 60-nt length in subsequent experiments to maintain consistency and maximize system performance.

Next, we evaluated the sensor’s ability to detect endogenous RNA transcripts. First, we tested whether SONAR could detect RNA expressed from a genomically integrated sequence introduced via the PiggyBac transposase system. For this, we generated a HEK293T cell line stably co-expressing the target RNA sequence and a MMLV-RT under an inducible promoter. qPCR confirmed that an efficient and specific RT-DNA synthesis occurred specifically in the presence of targeting ASOs (Supplementary Fig. 5a). Using the most effective ASO (Fig. 2c), we observed a significant 17.5-fold increase in SONAR activation for detecting the integrated target sequence compared to controls (Fig. 3a). Encouraged by these results, we further engineered sensors targeting different endogenous transcripts, including *EEF1A1* (Eukaryotic translation elongation factor 1 alpha 1), *GAPDH*, and *ACTB* mRNAs. For all transcripts, we designed and evaluated several ASOs targeting distinct regions within the 3′ UTR. All tested ASOs produced varying yet significant sensor activation compared to the non-targeting sensor control (Supplementary Fig. 5b–d). Subsequent validation with the top-performing ASOs, compared to stringent negative controls (including non-targeting ASO and no-ASO conditions), consistently induced strong and specific sensor activation, confirming SONAR’s specificity and reliability in detecting endogenous transcripts (Fig. 3b-d).

**Figure 3.**
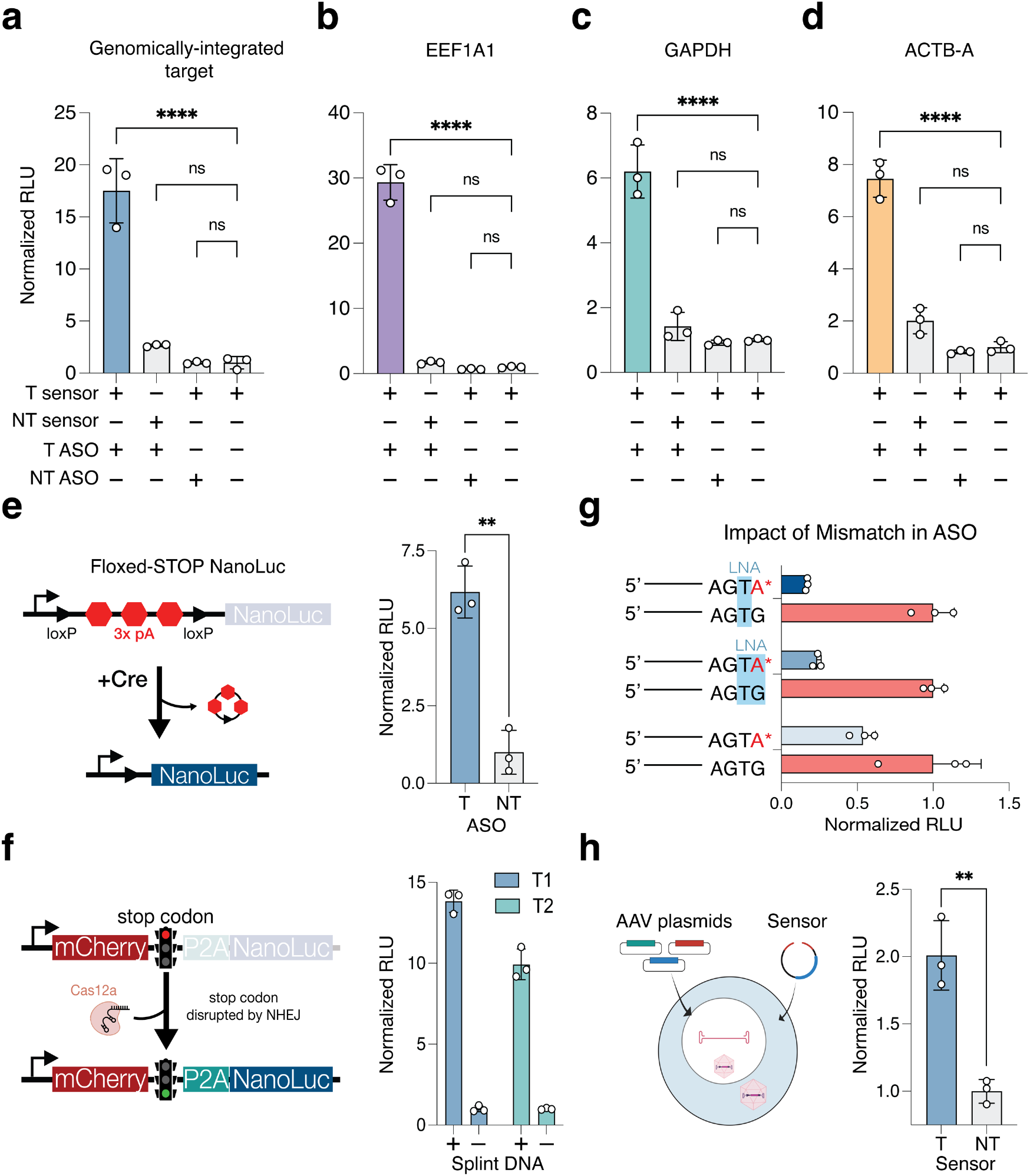
Detection of endogenous mRNA and SONAR applications. **a**. Quantification of sensor efficiency in detecting target RNA expressed from genomically integrated vectors in human cells. **b**. Quantification of sensor efficiency in detecting endogenous target RNAs: EEF1A1 **(b)**, GAPDH **(c)**, and ACTB **(d)** in human cells. For **a, b, c**, and **d**, the data was collected 48 h after SONAR transfection. Data are presented as fold change relative to the corresponding − ASO control and represent the mean of *n*=3 biological replicates. Significance determined by ordinary one-way ANOVA between all groups and − ASO control. \*\*\*\**P* < 0.0001, ns: not significant, *P* > 0.05. **e**. (Left) Schematic of the floxed-STOP luciferase reporter plasmid, in which three poly(A) signals flanked by loxP sites block luciferase expression until Cre-mediated excision. (Right) Quantification of sensor-driven Cre expression in detecting genomically integrated target RNA in human cells, measured as luciferase activity. The data collected 24 h after SONAR transfection. The data are presented as fold change of sensor with targeting ASO (T) relative to the corresponding non-targeting (NT) ASO control. Data represent the mean of n=3 biological replicates. Significance determined by unpaired two-tailed *t*-test.\*\**P* = 0.0012. **f**. (Left) Schematic of the Cas12a reporter plasmid, where a stop codon prevents downstream Nluc expression. Upon Cas12a and guide RNA (gRNA) expression, targeted cutting near the stop codon followed by non-homologous end joining (NHEJ) repair can restore the Nluc reading frame. (Right) Quantification of sensor efficiency in cells with or without the specific splint DNA. Data represent the mean of n=3 biological replicates. **g**. Impact of mismatches and LNA modifications on sensing specificity. Mismatches highlighted in red with an asterisk. LNA modifications are highlighted in light blue. Data represent sensor efficiency with ASOs containing mismatches relative to the corresponding perfectly matched ASOs. The data collected 24 h after SONAR transfection. Data represent the mean of n= 3 biological replicates. **h**. Quantification of sensor efficiency in detecting AAV genomic sequences in human cells. Data are from n=3 biological replicates. Significance determined by unpaired two-tailed *t*-test between targeting (T) and non-targeting (NT) sensor control. \*\**P* = 0.0031.

Some target sequences, e.g., for ACTB detection, inherently contain ATG codons, which could introduce unintended translation initiation at these sites, potentially disrupting proper SONAR-mediated payload expression. To address this challenge, we strategically introduced mismatches within the sensor sequences. Specifically, selected bases within ATG motifs were substituted, such as converting “T” to “G” or “A” to “C”. To verify that these intentional mismatches do not compromise sensor efficacy, we tested a sensor (ACTB-B) containing three such mismatches. Our results confirmed that this modified sensor maintained effective and highly specific detection of ACTB transcripts (Supplementary Fig. 5e and f). Notably, another sensor (ACTB-A), previously evaluated in Fig. 3d, also contained a single mismatch and similarly maintained high detection activity, demonstrating that targeted mismatch incorporation effectively mitigates unintended translation initiation without negatively impacting sensor performance. Together, these experiments highlight the flexibility, specificity, and robustness of the SONAR platform in detecting various endogenous transcripts, underscoring its potential for versatile, targeted RNA sensing across diverse biological applications.

As previously discussed, SONAR has the potential to function beyond a mere sensor, as it can trigger gene expression of any payload upon detecting RNA. This capability expands its use to more advanced gene regulation applications, such as driving programmable genetic circuits in human cells. To test this, we designed a system in which SONAR mediates the expression of a Cre recombinase, which in turn enables the expression of a luciferase reporter. We employed a floxed-stop reporter plasmid in which three polyadenylation (polyA) signals, flanked by Cre loxP sites, were inserted between a CMV promoter and an Nluc luciferase coding sequence, preventing luciferase expression without a Cre-mediated excision of the polyA signals (Fig. 3e). Initially, we validated that the Cre recombinase could be expressed from a circularizable ssDNA sensor upon the addition of a DNA splint, and that this Cre expression mediated a robust activation of the Nluc reporter (Supplementary Fig. 6a). We then tested whether this Cre-expressing SONAR system can be activated via RNA targets. The Cre SONAR system successfully detected target RNA from both a genomically integrated sequence and a plasmid-expressed target, resulting in a more than sixfold increase in luciferase activity compared to controls (Fig. 3e and Supplementary Fig. 6b). These findings highlight SONAR’s ability to function as a programmable genetic circuit, integrating RNA sensing with precise gene regulatory outputs.

To assess the capacity of the ssDNA sensor to express large genetic payloads while retaining its target-specific responsiveness, we engineered a sensor encoding Cas12a, a large genome-editing enzyme (>1300 amino acids)^17^. To validate Cas12a expression, we employed a reporter system consisting of an mCherry sequence followed by a stop codon and an out-of-frame Nluc sequence. Successful Cas12a expression and cleavage of the target sequence, i.e., the region containing the stop codon, can restore Nluc expression via a NHEJ repair-mediated frame shift (Fig. 3f). The functionality of this system was confirmed by co-transfecting a Cas12a-expressing plasmid (Supplementary Fig. 6c). Upon co-delivering the Cas12a-expressing ssDNA sensor and the reporter plasmid, we observed robust Nluc activation exclusively in the presence of the specific splint DNA, demonstrating the sensor’s capability to efficiently express of large proteins in response to specific nucleic acid sequences (Fig. 3f).

Accurate discrimination of SNPs is critical for biosensors, particularly in applications where distinguishing pathogenic mutations from benign sequences is essential. To address this challenge, we explored whether introducing synthetic mismatches on ASOs could enhance SONAR’s sensitivity and ability to differentiate SNPs. To test this, we designed a series of ASOs with strategic mismatches at different positions near the 3′ end and evaluated their impact on sensor activation. By comparing the sensor’s activation induced by each mismatched ASO to that of its perfect-match counterpart, we quantified SONAR’s sensitivity to single-base changes. Notably, the strongest fold difference (approximately 6-fold) was observed with an ASO carrying an LNA modification at the nucleotide preceding the 3′ terminus, or LNA 3′-1 (Fig. 3g and Supplementary Fig. 6d). Notably, such LNA modifications near the 3′ end also slightly reduced overall efficiency compared to ASOs without LNA modifications in these positions (Supplementary Fig. 6d). This suggests that ASOs might need to be optimized for each target SNP to ensure high specificity while maintaining sufficient efficiency. Overall, these findings demonstrate SONAR’s ability to distinguish SNPs, providing a promising approach for applications requiring high specificity in mutation detection.

Finally, we explored whether SONAR could detect viral DNA sequences in living cells, a capability that could be valuable for applications such as viral diagnostics, monitoring viral replication, and studying viral-host interactions. Unlike RNA detection, where reverse transcription and ASOs are required, direct detection of viral ssDNA genomes require the use of the ssDNA sensor only. To assess this, we transfected HEK293T cells with three plasmids required for adeno-associated virus (AAV) production, including one plasmid encoding the viral payload, i.e., the target sequence of interest. This process leads to the intracellular production of AAV ssDNA genomes. Employing an ssDNA sensor designed to recognize the target sequence within the AAV genome, we observed robust and sequence-specific detection of the viral DNA (Fig. 3h and Supplementary Fig. 6e). These findings demonstrate SONAR’s feasibility to detect viral genomes within living human cells.

## Discussion

Our findings demonstrate SONAR’s robust capability to specifically detect both exogenous and endogenous RNA targets with minimal optimization. Screening only a small number of ASOs allowed rapid identification of effective sensors, suggesting that further exploration could yield even more powerful designs. Additionally, our system consistently and reliably detected a diverse range of targets, highlighting its simplicity, and robustness across various experimental contexts. Furthermore, we showed that SONAR can extend beyond its sensing capabilities to drive the expression of effector proteins, such as Cre and Cas12a, in programmable genetic circuits.

SONAR successfully detects RNA targets by reverse-transcribing them into DNA via an exogenously provided RT. In future directions, this system could be further optimized by leveraging endogenous RTs, such as LINE1-^18^, Polθ-^19^, or telomerase^20^-derived reverse transcriptases. These RTs could be recruited via engineered ASOs or click chemistry, which would eliminate the dependency on an exogenous RT. Given SONAR’s robust performance in HEK293 cells, expanding this platform into other therapeutically relevant and primary cell types presents an exciting opportunity to enhance its clinical significance. This potential is particularly compelling since key components of SONAR, such as ASOs and RT, are rapidly progressing toward clinical translation^21,22^.

Finally, the demonstrated ability of SONAR to detect viral genomes within cells highlights a particularly intriguing therapeutic potential. Given the rising interest in ASOs as antiviral treatments^23–25^, SONAR could be envisioned in a dual-function-approach: using ASOs simultaneously as antiviral agents and as precise targeting modules to activate specific therapeutic responses within infected cells. Such integration would enable real-time detection and programmable cellular responses, significantly enhancing antiviral strategies.

By combining RNA and DNA detection capabilities, SONAR offers broad applications, including detecting pathogenic mutations, monitoring viral replication, and triggering precise gene expression tailored to individual molecular profiles. Collectively, these capabilities establish SONAR as a powerful, versatile platform poised to significantly advance precision diagnostics, targeted therapeutics, and fundamental biomedical research.

## Acknowledgments

G.M.C., A.M., L.M.R., and R.F. were supported by the Aging and Longevity-Related Research Fund. A.M was supported by the Ibn Rushd Postdoctoral Fellowship from King Abdullah University of Science and Technology.

## Author contributions

Conceptualized and initiated the study: A.M and R.F. Performed experiments: A.M., with help from R.F. and L.M.R. Analyzed the data: A.M. and R.F.

Supervised the project: A.M. and G.M.C.

Wrote the manuscript: A.M. and R.F. with input from all authors

## Methods

### Single-stranded DNA generation

All ssDNA constructs were synthesized by Genscript (GenExact™ Single-Stranded DNA Synthesis Service) with 5’ end phosphorylation. Each ssDNA molecule comprises a promoter (e.g., CMV) to drive transcription, followed by the sensing region divided into two segments flanking a nick site between the 3′ end of the upstream sensor and the 5′ end of the downstream sensor. Downstream of the sensing region is the gene of interest (GOI), which allows for the monitoring and quantification of gene expression. The construct is terminated with a PolyA sequence. The finalized ssDNA design is schematically represented as follows:

5′ –– downstream sensor - GOI - PolyA - CMV promoter - upstream sensor –– 3′.

All ssDNA sensors are listed in the supplementary file.

### Antisense oligonucleotides

All ASOs used in this study were obtained from Integrated DNA Technologies (IDT) as Affinity Plus antisense oligos. ASOs contained Locked Nucleic Acid (LNA) modifications at various nucleotides along the backbone, with the exception of the last four nucleotides at the 3’ end, which remained unmodified (Supplementary Fig.2c). Additionally, the first two or three nucleotides at the 5’ end of all ASOs were modified with phosphorothioate linkages. Other ASOs used for mutation detection had different patterns of modifications. All ASOs and modifications are listed in the supplementary file.

### Reverse transcriptase cloning

All plasmids were generated using standard molecular cloning methods, including Gibson assembly and restriction–ligation cloning. MMLV-RT plasmids were cloned by Gibson assembly of PCR products. Full-length MMLV-RT with nuclear localization signal (NLS) or lacking NLS were amplified from Addgene plasmid #190105 using Q5 Hot Start High-Fidelity 2X Master Mix (NEB). PCR products were purified using the Monarch Gel Extraction Kit (NEB), and then assembled under the control of the EF1α promoter into the backbone of Addgene plasmid #138272 (digested with SmaI and NotI) using the NEB HiFi DNA Assembly Master Mix.

To generate the RNase H-deficient MMLV-RT construct, the previously described MMLV-RT plasmid lacking NLS was PCR-amplified using primers designed to exclude the RNase H domain. The amplified PCR product was purified and then circularized by Gibson assembly with a short, double-stranded linker fragment, generated by annealing two complementary oligonucleotides (IDT).

### Target sequences cloning

Exogenous RNA target sequences were expressed from a modified version of Addgene plasmid #191138. To introduce custom sequences, the 3′ UTR of the original construct was replaced with a synthetic sequence generated by annealing two complementary oligonucleotides (IDT). These oligos were first phosphorylated and annealed using T4 Polynucleotide Kinase (NEB), then ligated into the NheI-and ApaI-digested #191138 backbone using T4 DNA Ligase (NEB). For the no-target control plasmid, the 3′ UTR sequence of the above plasmid was replaced with part of the mCherry coding sequence. This mCherry fragment was PCR-amplified and then Gibson-assembled into the backbone, which had been digested with XhoI and ApaI.

### PiggyBac plasmids

To generate stable cell lines co-expressing a target sequence and MMLV-RT, two PiggyBac (PB) transposon plasmids were constructed: (1) Target sequence PB plasmid: The target sequence was PCR-amplified from the previously described plasmid and cloned into the pB-CAGGS-dCas9-KRAB-MeCP2 vector (Addgene #110824), which carries a blasticidin resistance marker. The vector was digested with NheI and PmeI, and the insert was assembled using Gibson assembly. (2) MMLV-RT PB plasmid: The MMLV-RT coding region was PCR-amplified from the cloned MMLV-RT plasmid and inserted into a modified version of PB-TRE-dCas9-VPR (Addgene #63800), in which the hygromycin resistance cassette was replaced with puromycin. The recipient plasmid was digested with NheI and PmeI, and assembly was performed using Gibson assembly. This construct places MMLV-RT under the control of a TRE3G inducible promoter.

### Reporter plasmids cloning

For the floxed luciferase reporter plasmid for evaluating SONAR activity with a Cre payload, we utilized the pcDNA3.1_Floxed-STOP_Fluc plasmid (Addgene #122962), which expresses firefly luciferase (Fluc) under the control of a CMV promoter. This construct contains three polyadenylation (polyA) sequences flanked by loxP sites positioned between the CMV promoter and the Fluc coding sequence, thereby blocking downstream expression until Cre-mediated excision. To replace Fluc with Nluc, the Nluc coding sequence was PCR-amplified and cloned into the same vector digested with EcoRI and XhoI, generating the pcDNA3.1_Floxed-STOP_Nluc construct.

To construct Nluc reporter plasmids for evaluating SONAR activity with a Cas12 payload, synthetic gBlocks (IDT) encoding an mCherry sequence, a Cas12 target site, and an out-of-frame Nluc coding sequence were designed and cloned downstream of the CMV promoter into the plasmid backbone (Addgene #182540), which had been digested with NheI and XbaI, using the NEB HiFi DNA Assembly Master Mix. To construct Cas12 crRNA expression cassettes, synthetic gBlocks (Twist Bioscience) containing the U6 promoter and Cas12 crRNA sequences were cloned into Cas12 reporter plasmids linearized with MfeI using the NEB HiFi DNA Assembly Master Mix. Each U6-Cas12 crRNA cassette was integrated into its corresponding reporter plasmid, which harbored the Cas12 target site positioned between the mCherry and Nluc coding sequences. To evaluate the reporter functionality, enAsCas12a plasmid (Addgene#107941) was used.

### Tissue Culture

Cell lines were sourced from the American Type Culture Collection (ATCC, Manassas, Virginia) and maintained at 37 °C in and 5% CO2. The cells were cultured in Gibco™ DMEM with high glucose and pyruvate (ThermoFisher catalog #11-995-073), supplemented with 10% fetal bovine serum (FBS, ThermoFisher catalog #A5209501) and 1% Penicillin-Streptomycin (ThermoFisher catalog #15140122).

### Transient transfection

For all experiments, unless otherwise specified, approximately 2.5 × 104 HEK293T cells were seeded per well in 96-well tissue culture-treated plates and cultured to ~70% confluency prior to transfection. Transfections were performed using Lipofectamine 3000 (Invitrogen, L3000015). For each well, nucleic acid–Lipofectamine complexes were prepared by combining the desired plasmids or oligonucleotides with 0.35 µL of Lipofectamine 3000 reagent and 0.2 µL of P3000 enhancer in a total volume of 10 µL Opti-MEM (Thermo Fisher Scientific). Complexes were incubated for 20 minutes at room temperature before being added to the cells.

To assay for ssDNA sensor activation using splint sequences, cells were transfected with 25 ng of ssDNA sensor with GFP payload and 125 nM of specific DNA splint, non-specific DNA splint, or specific RNA splint. All oligos were purchased from (IDT).

For qPCR analysis of targeted ASO-mediated reverse transcription of endogenous RNA, HEK293T cells were grown in 24-well tissue culture-treated plates to ensure ample nucleic acid availability for subsequent analyses, and allowed to grow to approximately 70% confluency prior to transfection. The cells were first transfected with plasmids expressing reverse transcriptase. After 24 hours, the media was changed, and the cells were transfected with ASOs at a final concentration of 100 nM.

For detection of RNA transcripts expressed from plasmids, cells were first transfected with 40 ng of RT-expressing plasmids and 40 ng of plasmid encoding the RNA target sequence, or a non-targeting control plasmid as a negative control. After 24 hours, the media was changed, and cells were transfected with 25 ng of ssDNA sensor along with the corresponding antisense oligonucleotides (ASOs). Unless otherwise specified, ASOs were used at a final concentration of 125–150 nM in all experiments involving plasmid-expressed RNA targets.

For detection of endogenous RNA transcripts, cells were first transfected with 50 ng of RT-expressing plasmids. After 24 hours, the media was changed, and the cells were transfected with 25 ng of ssDNA sensor mixed with 125 nM of the respective ASOs.

For detection of a genomically integrated target, approximately 2.8 × 104 HEK293T cells stably expressing the target sequence and an inducible MMLV-RT were seeded into 96-well tissue culture-treated plates in media supplemented with 1.5 µg/mL doxycycline. After 24 hours, once the cells reached approximately 70% confluency, they were transfected with 25 ng of ssDNA sensor and 125 nM of the corresponding ASOs.

For the floxed luciferase assay with Cre-SONAR, cells were first transfected with 40 ng of RT-expressing plasmids and 40 ng of plasmid encoding the RNA target sequence, or a non-targeting control plasmid as a negative control. After 24 hours, the media was changed, and cells were transfected with 5 ng of pcDNA3.1_Floxed-STOP_Nluc reporter, 5 ng of the ssDNA sensor encoding Cre recombinase, and 200 nM of the corresponding ASO. On day 3, the cells were processed following the protocol described in the “Luciferase Assays” section below.

For the floxed luciferase assay with Cre-SONAR targeting a genomically integrated target, HEK293T cells expressing the target sequence with inducible RT were prepared as discussed above. When the cells reached approximately 70% confluency prior to transfection (24 hours), they were transfected with 5 ng of pcDNA3.1_Floxed-STOP_Nluc reporter, 5 ng of the ssDNA sensor encoding Cre recombinase, and 200 nM of the corresponding ASO. After 24 hours, the cells were processed following the protocol described in the Luciferase Assays section.

For Cas12 detection assays using plasmid-expressed enAsCas12a, cells were transfected with 10 ng of enAsCas12a plasmid and 20 ng of the Nluc Cas12 reporter plasmid, which served both as a surrogate reporter for Cas12 activity and as an expression cassette for the Cas12 crRNA. For the no-Cas12 control, 10 ng of a non-targeting plasmid was used in place of the Cas12 plasmid. After 48 hours, cells were processed according to the protocol outlined in the Luciferase Assays section.

For Cas12-SONAR detection assays, cells were transfected with 25 ng of Cas12 ssDNA sensor, 10 ng of Nluc Cas12 reporters, and either 125 nM or 0 nM of specific DNA splint. After 48 hours, the cells were processed following the protocol described in the Luciferase Assays section.

For detection of AAV genomic ssDNA using the sensor, HEK293T cells were seeded in 96-well tissue culture-treated plates. Once the cells reached approximately 50% confluency, they were transfected with AAV packaging plasmids, including 38 ng of a cis-plasmid containing an ITR-flanked BFP-encoding sequence with the sensor target site under the EF1A promoter (VectorBuilder), 38 ng of a Rep/Cap plasmid (Addgene #104963), and 45 ng of an AAV helper plasmid (Addgene #112867). After 34 hours, the media was changed, and cells were transfected with 50 ng of ssDNA sensor. 24h after sensor transfection, cells were processed according to the protocol described in the Luciferase Assays section.

### Cell line generation

HEK293T cells were seeded in 6-well tissue culture-treated plates and cultured until approximately 75% confluency before transfection. Cells were transfected with 2.5 µg of the cloned piggyBac plasmids (as described above) and 1 µg of Super piggyBac Transposase plasmid (System Biosciences, PB210PA-1) using Lipofectamine 3000. Forty-eight hours post-transfection, selection was initiated by adding 2 µg/mL puromycin and 5 µg/mL blasticidin to the culture medium, depending on the resistance markers encoded by the plasmids. Cells were maintained under these selective conditions for two weeks to ensure stable integration and selection of transfected cells.

### Luciferase assays

To measure luciferase activity, we used the NanoLuc (Nluc) luciferase reporter system (Promega). At 24 or 48 hours post-transfection (as specified in the figure legends), cells were washed with phosphate-buffered saline (PBS), and the Nano-Glo Luciferase Assay (Promega) was applied directly to the cells according to the manufacturer’s instructions. Lysates were transferred to black 96-well plates with clear bottoms (Costar 3603, Corning), and luminescence signals were measured using a BioTek Synergy H1 microplate reader.

### Fluorescence Microscopy

Fluorescent signals within the cells were detected and analyzed using the FLoid™ Cell Imaging Station (Life Technologies).

### Quantitative PCR Analysis

Twelve hours after ASO transfection, both RNA and ssDNA were extracted using the RNeasy Plus Mini Kit (QIAGEN, 74134). Following extraction, samples were treated with RNaseA (NEB, T3018) for 15 mins and followed by ezDNase treatment (Invitrogen, 117660512) for 4 mins. ssDNA was then purified and concentrated using Monarch® Spin PCR & DNA Cleanup Kit (NEB, T1130). 2 µL of each sample was prepared for quantitative PCR (qPCR) using the Luna® Universal qPCR Master Mix (New England Biolabs). qPCR was conducted on a Roche LightCycler® 96 platform with the following thermal cycling protocol: an initial denaturation at 95°C for 60 seconds, followed by 40 cycles of denaturation at 95°C for 15 seconds and extension at 60°C for 30 seconds, and culminating with a melt curve analysis from 60°C to 95°C. Cycle quantification (Cq) values were automatically calculated using the Roche LightCycler® 96 software. All qPCR reactions were performed in triplicate to ensure reproducibility and reliability of the results.

### PCR amplification of splint-ligated ssDNA sensor

Cells were transfected with varying concentrations of the ssDNA-GFP sensor, with or without the specific splint DNA, as described above. Genomic DNA was collected 24 hours post-transfection by removing the medium, washing the cells once with PBS, and resuspending them in 100 µL of QuickExtract™ DNA Extraction Solution (Biosearch Technologies, QE09050). The lysate was incubated at 66 °C for 6 minutes followed by 98 °C for 2 minutes to complete the extraction. Subsequently, 2.5 µL of the extracted genomic DNA was used as input in 50 µL PCR reactions using Q5 Hot Start High-Fidelity 2X Master Mix (NEB).

## Supplementary Figures

**Supplementary Figure 1.**
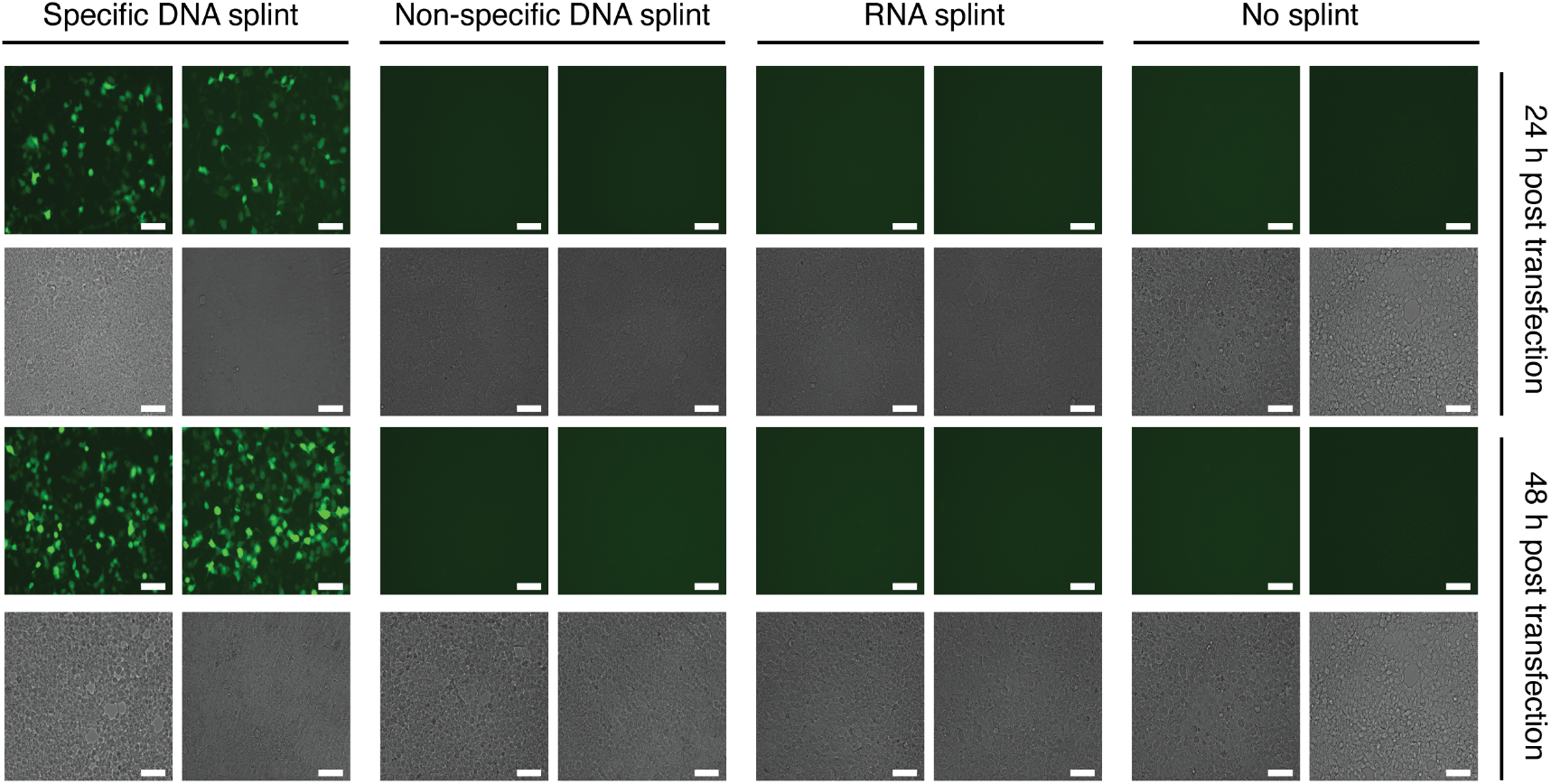
Evaluation of ssDNA sensor activity and specificity. Fluorescence images of HEK293T cells transfected with the ssDNA sensor and either a complementary DNA splint or control conditions after 24h (top) and 48h (bottom). Scale bar: 125 µm.

**Supplementary Figure 2.**
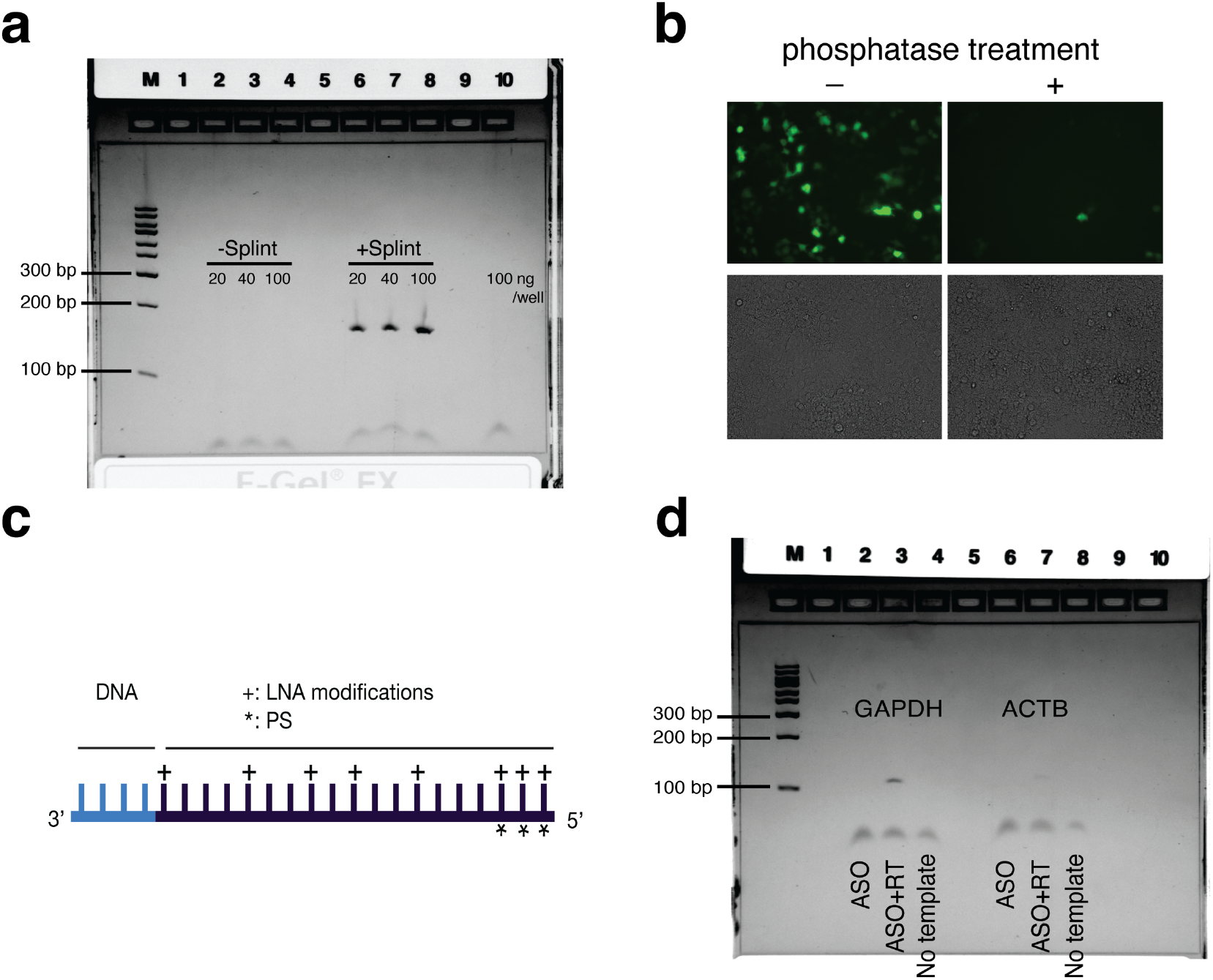
Characterization of ssDNA sensor. **a**. Uncropped gel electrophoresis of PCR-amplified sensor described in Fig.1c. **b**. Impact of phosphatase treatment on the sensor’s activity. **c**. Schematic of the ASOs used in the study. **d**. Uncropped gel electrophoresis of PCR product of GAPDH and ACTB experiments described in Fig.2a.

**Supplementary Figure 3.**
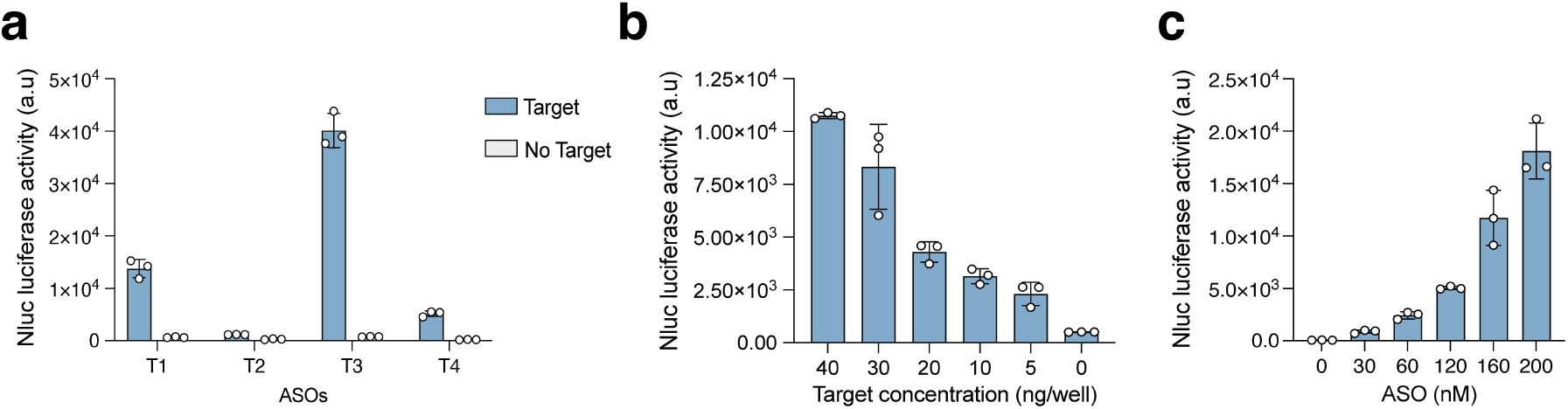
Raw data of sensor efficiency in detecting target RNA in human cells (from Fig.2c, d, and e). Data are means of *n*=3 biological replicates.

**Supplementary Figure 4.**
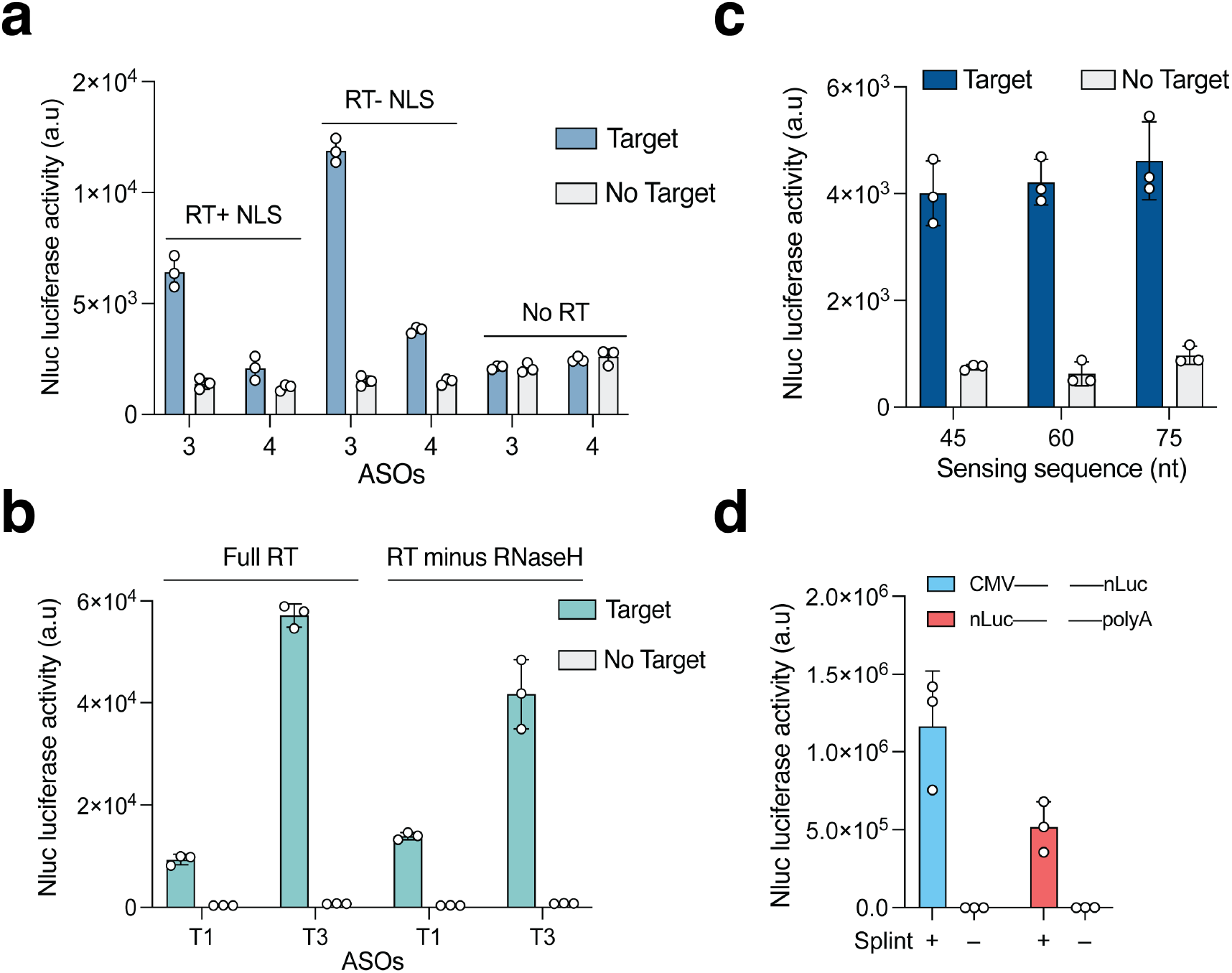
Activity of different reverse transcriptases, length and positioning of the sensing sequence. **a**. Evaluation of the SONAR efficiency using RT with nuclear localization signal (RT+NLS), without NLS (RT-NLS), or with No RT. **b**. Evaluation of the SONAR efficiency using RT with and without its RNase H domain. **c**. Impact of nucleotide length of the sensing sequence on SONAR activation. **f**. Quantification of SONAR activity with two different ssDNA sensor designs. CMV--Nluc: ssDNA sensor with the sensing region placed between the promoter and the Nluc payload sequence. Nluc--PolyA: ssDNA sensor with the sensing region placed between the Nluc payload sequence and the poly(A) tail sequence. In **a, b, c**, and **d**, data represent the mean of *n*=3 biological replicates.

**Supplementary Figure 5.**
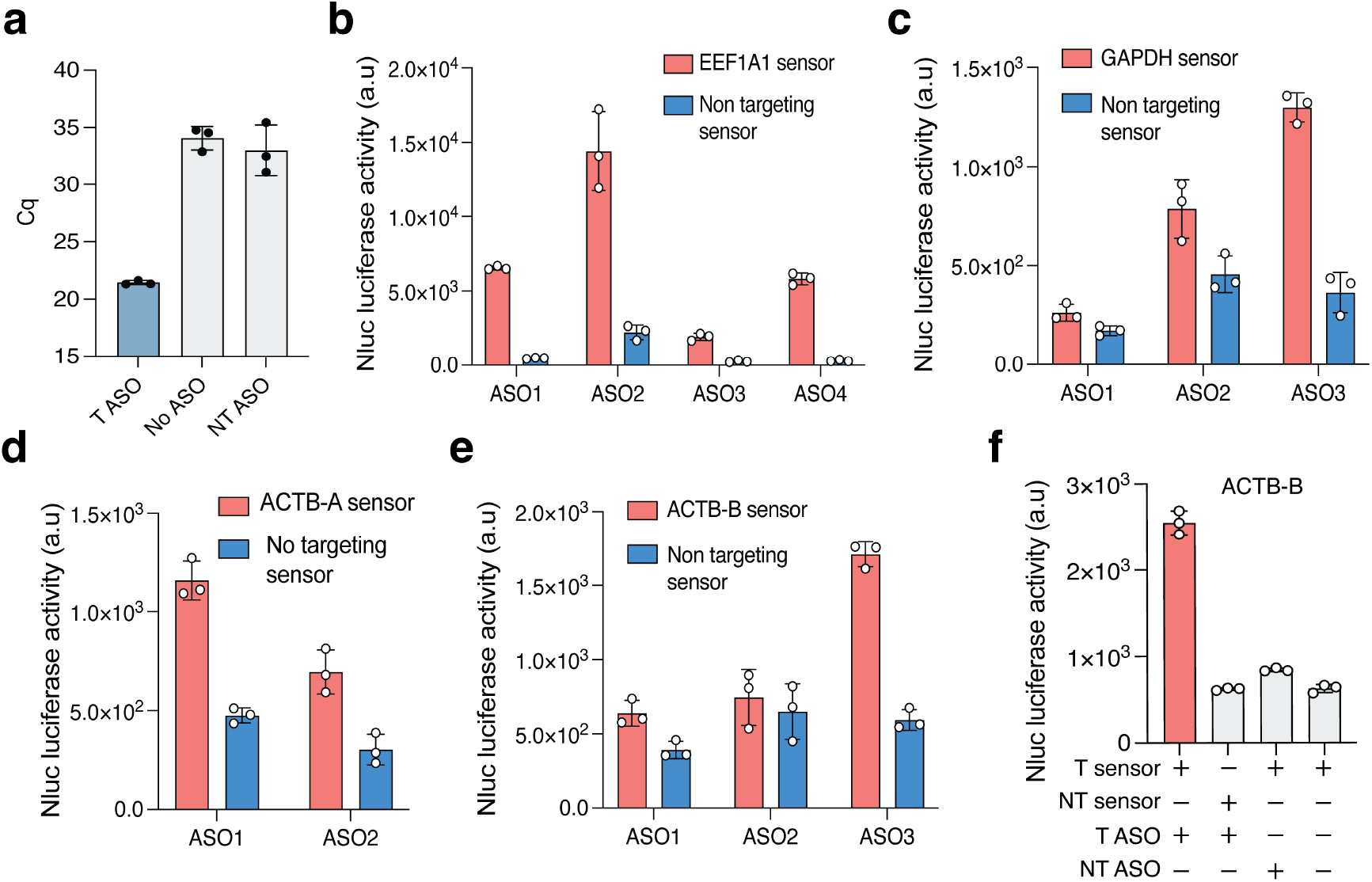
Raw data of SONAR activity for detection of endogenous transcripts. **a**. Quantitative PCR (qPCR) analysis of in-cell-synthesized reverse-transcribed DNA (RT-DNA) from a genomically integrated target. T ASO: targeting ASO, NT ASO: non targeting ASO. **b**. Raw luciferase values for screening ASO efficiency for *EEFA1A1, GAPDH* **(c)**, *ACTB* sensor A **(d)**, and *ACTB* sensor B **(e). f**. Extended control data for ACTB-B using ASO3, with raw luciferase measurements under various conditions. For **a, b, c, d, e**, and **f**, data represent the mean of *n*=3 biological replicates.

**Supplementary Figure 6.**
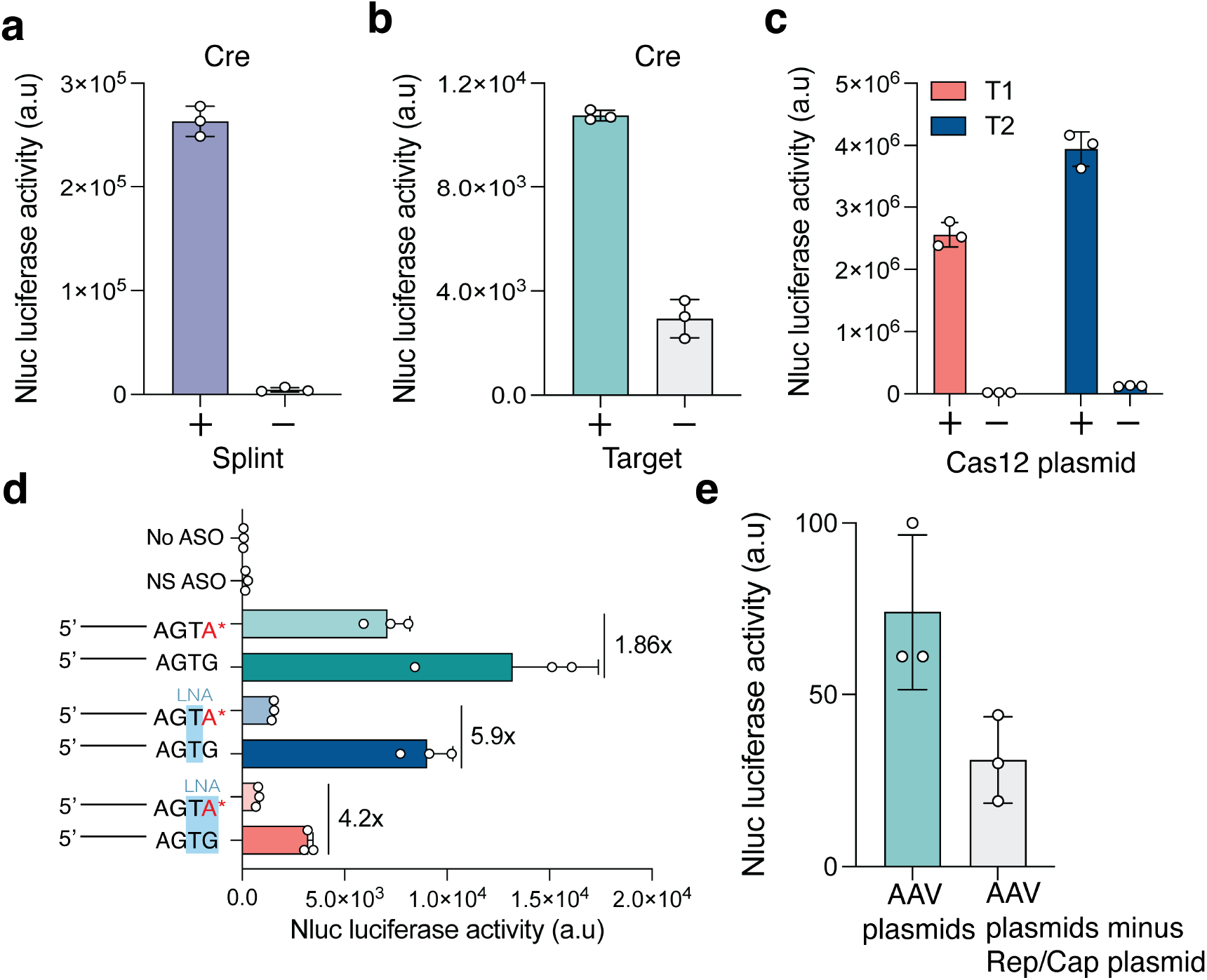
SONAR Applications. **a**. Assessment of the activity and target responsiveness of Cre-containing ssDNA sensor, evaluated through sensor-driven Cre expression in response to a specific splint DNA sequence in human cells. Activation was measured via Nluc reporter luciferase activity, with luminescence readings collected 24 hours post-transfection. Data represent the mean of *n*= 3 biological replicates. **b**. Quantification of sensor-driven Cre expression in detecting plasmid-expressed target RNA in human cells, measured as luciferase activity. The data collected 24 h after SONAR transfection. Data represent the mean of *n*=3 biological replicates. **c**. Assessment of the Cas12 Nluc reporter functionality using Cas12-expressing plasmid and two different reporter plasmids containing different Cas12 target sites (T1 and T2) and a Nluc sequence. Cas12-mediated cleavage of the reporter can lead to NHEJ-mediated repair, restoring the correct reading frame and resulting in Nluc expression. Luciferase activity was measured to evaluate Cas12-induced reporter activation with luminescence readings collected 48 hours post-transfection. Data represent the mean of *n*= 3 biological replicates. **d**. Raw values of data in (Fig. 3g) showing the impact of mismatches and LNA modifications on sensing specificity. Mismatches highlighted in red with an asterisk. LNA modifications are highlighted in light blue. The data collected 24 h after SONAR transfection. Data represent the mean of *n*= 3 biological replicates. **e**. Quantification of sensor efficiency in detecting AAV genomic sequences in human cells. AAV plasmids indicate the three AAV production plasmids needed for AAV replication, including transgenic gene (ITR) plasmid, Cap/Rep plasmid, and helper plasmid. AAV plasmids minus Rep/Cap control are cell cells transfected with AAV plasmid, but without Rep/Cap plasmid essential for AAV replications. Data are from *n*=3 biological replicates.

